# *Temnothorax rugatulus* ants do not change their nest walls in response to environmental humidity

**DOI:** 10.1101/2022.06.30.497551

**Authors:** Greg T. Chism, Wiley Faron, Anna Dornhaus

## Abstract

Animal architectures are interesting biological phenomena that can greatly increase the fitness of the builder and exist in a variety of forms and functions across taxa. Among the most intricate architectures are social insect nests, which may have several functions, one of which is the control of internal microclimate. In social insects, the regulation particularly of humidity in the nest can be crucial for the survival and growth of the brood. Though much is known on how nest *excavating* social insects respond to environmental humidity, little is known about how ants that build on to pre-existing cavities respond. Here we use the rock ant *Temnothorax rugatulus* to determine whether and how colonies respond to environmental humidity by building and changing their nest architectures in pre-existing nest spaces. We specifically test the hypothesis that *T. rugatulus* colonies build different nest walls, e.g. wider or denser ones, in response to lower environmental humidity. We allowed *T. rugatulus* colonies to build nest walls with two substrates across a 0-100% relative humidity gradient. We further compare the porosity - empty volume in built nest walls - of natural *T. rugatulus* nest walls with these artificial building substrates and the substrate compositions of built walls from our experiment. We found that humidity did not influence the nest walls *T. rugatulus* colonies built in our experiment, concluding that regulating humidity is likely not a key function of *T. rugatulus* nest wall architecture. We also found that the porosities of the artificial substrate that was predominantly used by the ants in our experiment were like the porosity of natural *T. rugatulus* nest walls, indicating that ants had constant preferences for particular substrates. Physical nest wall features, including porosity, are therefore unlikely to be flexibly regulated in response to external humidity, but may be adaptations in other ways.

## Introduction

Ants are some of the most ecologically diverse and evolutionarily successful organisms [1] likely at least in part due to the existence of nests that span in complexity from simple one-chambered spaces to complex spaces with several dozen chambers [2–8]. Nests serve as a stable and defensible space that likely facilitated the development of the social lifestyle and reproductive division of labor, i.e. a separation between reproducing individuals and non-reproducing ‘workers’ [1, 9]. Due to this intimate tie to their nest, social insect colonies have been referred to as a ‘factory within a fortress’ [9].

Many social insects modify their nests to respond to different environmental conditions to control their microclimate [10]. Regulating humidity for example has been shown to be a key role in the nest architecture in some social insects, such as leaf cutter ants, in which workers plug nest holes to prevent desiccation [11, 12] and mound building termites which construct a sophisticated nest ventilation system that constantly changes throughout the day [13-15] Humidity is particularly relevant to ants since larvae and eggs desiccate in dry conditions [16-18]. Though we know much about how humidity influences social insect colonies such as the above that excavate their nests by removing substrate from the ground, the effect of the abiotic environment on social insect colonies that build nests by adding material, such as walls in rock crevices (additive nest builders), are not well studied.

*Temnothorax rugatulus* ants are found in pine and juniper zones in northern Mexico, the western United States and southwestern Canada, and have a colony size between 50 to 400 ants [19]. They reside in preexisting structures such as rock crevices and acorns (i.e. arboreal and hypogaeic), thus creating a dark, cool, and humid nest space. Members of the *Temnothorax* genus are additive builders that utilize stones (i.e. what might, at this scale, be called sand grains) and other environmental substrates to produce walls that change their occupied nest space [20-21]. *Temnothorax* colonies choose a smaller-grained substrate when carrying is less energetically expensive (e.g. placed closer to the nest) [22-23], but larger stones are preferred during nest expansion and contraction [24], possibly because of the need for speed in building. *Temnothorax rugatulus* colonies both build thicker walls when they have more brood, and build longer, larger area walls in higher environmental humidity [25]. However, in that study, colonies were only offered one grain type. Walls built with two stone sizes that are well mixed produce a higher angle stability and thus are more structurally stable, because smaller grains can fill in the gaps of larger grains [23]. We propose that mixed walls may also regulate nest microclimate more efficiently than single substrate walls by creating a more compact structure.

In this study we asked whether the ant *Temnothorax rugatulus* produces different nest walls in response to differing environmental humidity. Response to humidity is reasonable since *T. rugatulus* colonies live inside crevices found in granite boulders in a desert environment that likely experience occasional high ground temperatures and low environmental humidity. In this study, we had *Temnothorax rugatulus* build nests under different relative humidity levels, and then quantified differences in nest properties. We also tested whether the nest properties scaled with colony size. To confirm whether environmental humidity constitutes a selective force on these ants, we also measured worker and brood mortality under the humidity experienced in this experiment. Brood and workers are vulnerable to desiccation, so we would expect lower environmental humidity to cause more death. We also compared the porosity of *T. rugatulus* walls from our experiment and natural walls collected in the field, which may relate to the degree of moisture saturation that the nest wall may exhibit.

## Methods

### Colony collections

We collected twenty-two colonies of *Temnothorax rugatulus* from the Santa Catalina Mountains (GPS: 32.395, -110.688), USA, Pima County, Arizona in a pine-forest zone (altitude approximately 2500m) in October 2018. From February to October 2018, we further collected ten separate *T. rugatulus* complete nest walls (e.g. in Fig 1a), which were obtained after the entire colony was removed. We only collected unbroken walls to be certain that all substrate was accounted for.

**Fig 1.**
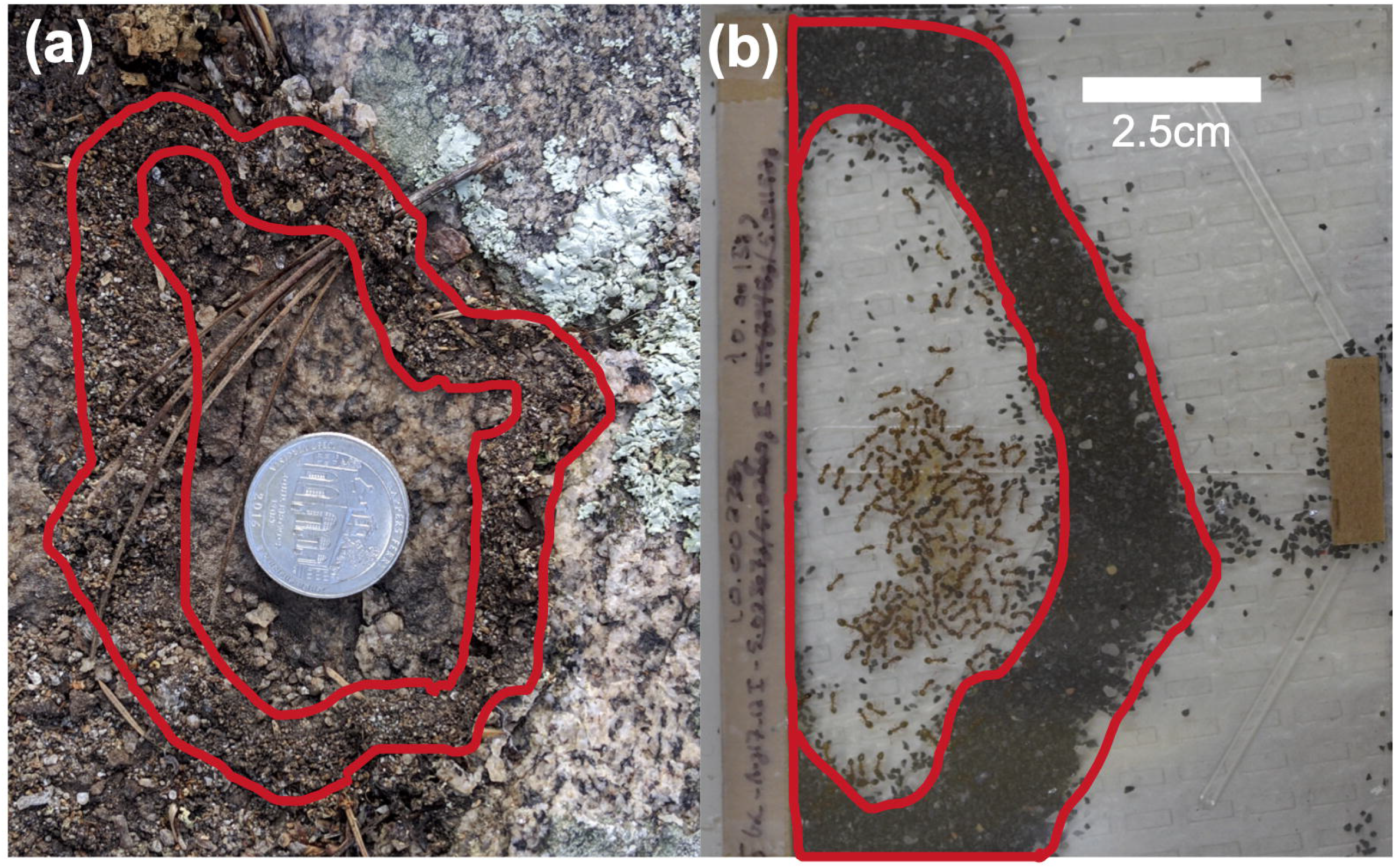
Comparing a natural nest wall of a *Temnothorax rugatulus* colony in the field (a) and wall built by a *T. rugatulus* colony experimentally (b). Notably, the natural nest wall is moist and caked together, where the stone (sand) walls in the lab can easily be pushed apart. In both photos, the outer and inner boundaries represent the built nest wall and inside of the inner boundary is where the queen(s), workers, and brood reside (internal nest area).

### Colony acclimation period

We first acclimated our experimental colonies in a controlled environment produced in a climate chamber for five days (see Initial housing and care). We determined experimental length through a preliminary building assay, where we allowed the colonies to build nests using experimental building substrates (see description below) for twenty days and found no substantial nest wall changes after 10 days. We therefore set 10 days as the building duration for our future experimental building phases.

#### Initial housing and care

During the acclimation period, we housed the colonies in 17.5cm x 12.5cm x 6cm plastic containers with inside walls coated in ‘insect-a-slip’ (BioQuip product #2871A) to prevent escape. We gave each colony a nest space made of two glass panes (102mm x 76mm) separated by a 1.5-mm-thick strip of cardboard at the back and smaller piece of cardboard at the front [25]. On the opposite end of the container, we gave each colony a water-filled 5 ml plastic tube with a cotton ball stopper and fed each colony *ad libitum* weekly with both a 2ml microcentrifuge tube of honey water with a concentration of 1/4 teaspoon per 50ml water, and 1/8 of a fresh-frozen cockroach (approximately 0.075g) (*Nauphoeta cinerea*). During acclimation and between experimental trials, we kept colonies in a climate chamber with a 12:12 h light cycle (8 a.m. to 8 p.m.), constant temperature (approximately 20°C) and relative humidity (approximately 20-25%).

### Experimental timeline

We exposed each colony to one of nine humidity levels (see S1 Table). We allowed the humidity in each container to stabilize for one day, at which point we then provided the experimental building substrates for colonies to build for 10 days. We photographed the colony on days 1, 5, and 10 to determine the average number of workers and brood, but only considered day 10 for calculating nest wall properties. We gave colonies a 10-day rest period in ambient temperature and humidity before the second trial in which we placed the colonies into the humidity three places forward along the gradient (i.e. 55% went to 85%, and 85% went to 1%).

### Experimental building substrates

We chose two distinct building substrates to allow *T. rugatulus* colonies to modify their nest spaces. We offered 10g of both 1.25mm diameter white aquarium substrate (CaribSea Super Naturals™ white aquarium substrate: substrate I - weight of 100 pieces = 0.529g) and 0.65mm diameter black aquarium substrate (Flouritel1l black sand: substrate II - weight of 100 pieces = 0.035g) as building materials. Though substrate I would cover more area per grain when building, its weight per grain is 15x larger than substrate II, making it harder to transport.

### Experimental setup

#### Experimental nest housing

We placed colonies in new nest sites and containers following the same protocol as during the acclimation period (see Fig 2 for a full set visualization).

**Fig 2.**
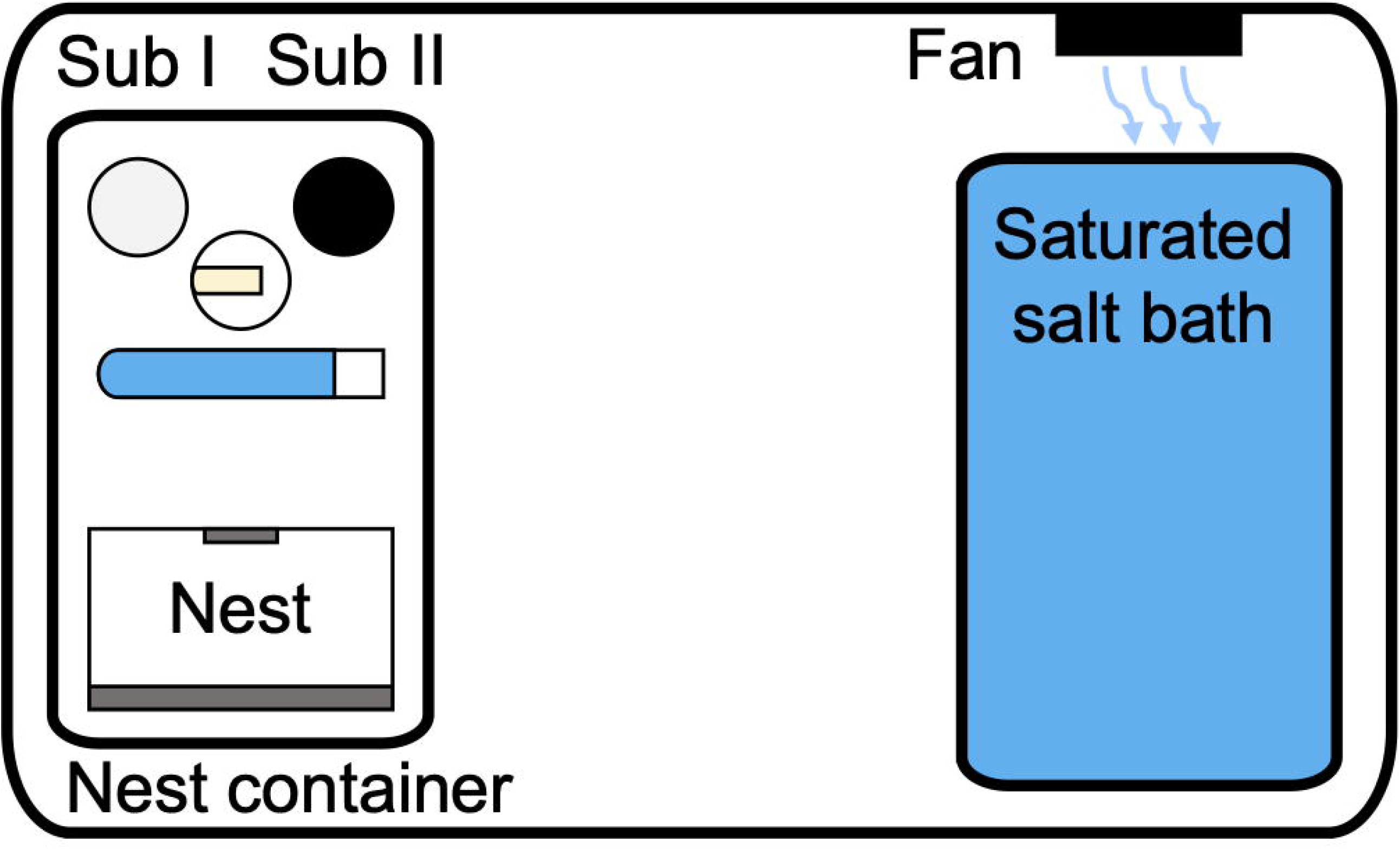
Experimental set up for each humidity treatment. Ants are confined to the nest container, which contains a small pile of each building substrate and food and water. A small fan gently circulated air above the saturated salt water to ensure homogenous humidity across the system. While this system was closed (locked airtight), a small hole was cut from the lid allowing access to the nest container to deposit each building substrate, thus temporarily opening the system when needed.

#### Experimental humidity levels

For each experimental trial, we placed the nine individual colonies and their nest containers into larger plastic containers (31.3cm x 23cm x 10.2cm). We created a discrete-step humidity gradient consisting of 9 separate boxes, each with a different, constant, regulated humidity level. Eight of these were achieved by using saturated salt solutions, which produce highly replicable humidity levels at 20°C in a closed, airtight system (Fig 2; S1 Table; Winston and Bates 1960; Greenspan 1977). We used the desiccant phosphorous pentoxide to produce nearly 0% RH (Winston and Bates 1960). We placed each saturated salt solution and desiccant in a plastic container (17.5cm x 12.5cm x 6cm) next to the nest container in the experimental setup (Fig 2). We used a DC current fan (4cm x 4cm x 4cm, 12V, 0.1A) in the top left corner of the experimental setup to circulate air in the closed system, since saturated salt solutions require air circulation for reproducibility (Winston and Bates 1960). This fan was placed in the top left corner above the saturated salt solution, angled horizontally, facing parallel to the length of the box (Fig 2).

#### Substrate, food, and water placement

We inserted substrates, food, and water through a small 3.5cm hole above the nest section, which only temporarily broke the closed, airtight system of the colony box. We placed individual 10.0g piles of each building substrate at the opposite end of the housing nest, such that one substrate type was on the left and the other on the right. On day 7 of the experiment, we provided new 5ml cotton-ball-clogged water tubes, 5ml honey water microcentrifuge tubes, and 1/8 fresh-frozen cockroaches to continue *ad libitum* feeding. We randomized the arrangement of the building substrates such that half of the colonies had the heavier substrate on the left side and the other half on the right side (Fig 2). During the second trial we flipped each colony’s substrate placement. We performed this procedure to prevent a side bias from affecting building substrate choice.

### Data collection

#### Environmental data

During each experimental round, we recorded the temperature (°C) and relative humidity (%) of our experimental closed systems every 45 minutes using permanently imbedded U12-012 HOBO data loggers (Onset, Bourne, MA, USA) to ensure stability and reproducibility of each section of the relative humidity gradient (S1 Table for experimentally produced humidity levels).

#### Image capture and analysis

We photographed each colony with an HD camera (Nikon D7000 with 60mm lens). We used the image analysis software *Fiji* [28] to measure the wall length (mm), wall area (mm^2^), and nest area (mm^2^) for every colony, which are measurement methods we derived from [25]. Additionally, we assigned coordinates to every worker and brood item in the nest. We standardized all measurements and coordinates from *Fiji* using the x-distance between the top left and bottom right of the glass pane as reference points as the known distance of 102mm.

#### Nest wall composition

After each building period, we collected each built wall by gently tilting a colony’s housing nest space and extracting the grains while the ants were inside the nest, allowing us to prevent panic. We sieved the nest walls built by each colony using a 1mm colander to separate the two substrates, and then weighed each substrate using a digital scale (Ohaus, USA) to the nearest 0.00001g.

#### Nest wall weight and density

We determined wall weight by weighing the substrates each colony used to build their nest walls. We calculated wall volume (mm^3^) as the nest area multiplied by 1.5 mm (the height of the provided nest cavity). We then calculated nest wall density as total wall weight (g) / wall volume (mm^3^).

#### Building substrate porosity

We used the collected substrates (see colony collections) of real *T. rugatulus* nest walls to compare the porosity between natural nests and our artificial building substrates. We allowed the substrates to dry in open air for seven days before storing them again. Our final sample size was 10 measures of porosity for the natural and each experimental substrate. Porosity (*Pt*) is calculated by determining the void space (*Vp*) in which water can fill in a substrate and dividing it by the bulk volume (*Vt*) which is the void and substrate (*Vs*) volumes: *Vt* = *Vp* + *Vs*; *Pt* = (*Vp*/*Vt*) x 100.

#### Experimental substrates

We measured the porosity of each artificial substrate by filling a 5 ml tube with 2 ml of each substrate (total volume: *Vt*). We determined the pore volume by fully saturating the substrate with deionized water injected through a syringe to the 2 ml mark. We took water from a container of water (weighed to the nearest 0.001g) and then determined the volume used (*Vp*) by subtracting the remaining container weight from the original weight of water. We converted water weight to volume per the 1g/ml standard conversion for pure water. We then calculated porosity for each substrate using the formula: *Pt* = (*Vp*/*Vt*) x 100.

#### Natural nest walls

To compare the porosities of natural and experimentally built nest walls, we measured the porosity of natural wall substrates by filling 5 ml tubes with each natural nest’s substrates and then marking where the tube was filled. We again filled the container with deionized water from a container of water (weighed to the nearest 0.001g) to complete saturation at the marked substrate volume line, then we subtracted the remaining container weight from the original weight of water to determine the pore volume (*Vp*). We removed the substrates from the containers and filled water to the line representing the volume of each substrate and weighed that amount to the nearest 0.001g (*Vt*). We again converted volume from water weight to volume per the standard 1g/ml conversion for pure water. We then calculated porosity for each substrate using the formula: *Pt* = (*Vp*/*Vt*) x 100.

### Final data and analyses

We conducted all data wrangling, analyses, and visualizations in the software R (*v4*.*1*.*1*) [29] in RStudio (*v1*.*2*.*5042*) [30], primarily utilizing the tidyverse language (‘tidyverse’ *v1*.*3*.*1*) [31]. We have made the final data and R script used for this study openly available in a GitHub repository: https://github.com/Gchism94/HumidityProject

#### Humidity treatment final data set

We had different final sample sizes for the first (trial 1) and second (trial 2) humidity treatments a colony experienced. We did not include three colonies in our final trial 1 data set due to colony death (trial 1 final N = 19 colonies). Three colonies were not placed into a second treatment since the first relative humidity produced in the first trial was not reliable (trial 2 final N = 16).

#### Humidity treatment analyses

Substrate preference: we used Mann-Whitney U tests to see whether colonies built their walls with a preferred substrate.

Influence of humidity and colony size on built walls: we took average values for worker and brood number (see image capture and analysis) to reduce measurement error from worker and brood occlusion in nest containers. We used linear mixed effects models using the packages to examine whether relative humidity or colony size influenced nest wall feature, or internal nest area. All linear mixed effects models here and below were conducted using the R package ‘lme4’ (*v1*.*1-27*.*1*; Bates et al., 2014), where p values were calculated through the R package ‘lmerTest’ (*v3*.*1-3*; Kuznetsova et al., 2017). We found that an order effect was present when considering a colony’s first and second trial placement, which we could not separate from the humidity treatment placement due to unequal sample sizes (lower:higher N = 14; higher:lower = 5). We therefore assigned the first and second trial (‘Trial’) as a random effect in our models, and by comparing the variation explained by the fixed effects alone (marginal R^2^) and with the random effect included (conditional R^2^), we determined the amount of variation that Trial number explained (marginal and conditional R^2^ values calculated through the R package ‘MuMIn’ *v1*.*43*.*17*; Kamil Bartoń 2020).

Worker and brood mortality and humidity exposure: we first used a linear regression to test whether brood and worker death were correlated. We then calculated worker and brood mortality as the proportion of average workers or brood in trial 2 over the average in trial 1, signifying how many of each died in between trials (10 days). We used binomial family generalized linear models to test whether the relative mortality of workers and brood was affected by the level of relative humidity each colony experienced in trial 1. Finally, we tested whether colony mortality was higher in smaller or larger colonies (average number of brood or workers) by comparing the two linear regressions with an ANOVA: formula = log(Trial 2 colony size) ∼ log(Trial 1 colony size); formula = log(Trial 2 colony size) ∼ 1 + offset(Trial 1 colony size). The offset function changes the model intercept to 1, where smaller colonies experienced higher mortality with a model intercept smaller than 1 and larger colonies experienced higher mortality with a model intercept greater than 1.

#### Building substrate porosity analysis

We used pairwise Dunn’s tests with False Discovery Rate corrected p values [35] to compare the median porosity of our experimental and collected natural nest wall substrates (each N = 10).

#### Post hoc power analyses

We used the R package simr (*v1*.*0*.*6*) [36, 37] to calculate post hoc power analyses for each of our linear mixed effects models where we used 0.5 and 0.8 as biologically relevant Cohen’s d effect sizes (moderate and high effect sizes) [38]. We first derived the appropriate effect size for each model (‘Humidity’ fixed effect β coefficient) from the formulas: Cohen’s d = β */* (sqrt(N) x SE), where N = sample size and SE = standard error of each ‘Humidity’ term. The simr package determines statistical power by (i) simulating new values for the response variable from our models; (ii) refitting our models to the simulated response variable; (iii) applying a likelihood ratio test to the simulated model fit. Statistical power is then determined by the ratio of significant p values over non-significant.

## Results

### Ants prefer smaller-grained substrate but show no side bias

We used two-sided Wilcoxon tests with a predicted median value of 0.5 to show that ants showed a substrate preference in trial 1 (W = 183, p < 0.001) and trial 2 (W = 132, p < 0.001). The predominant substrate that ants used for building was the smaller-grained substrate: in trial 1, the median weight of substrate I that colonies used to build walls 0.160g, and 0.516g for substrate II, with the median proportion of substrate II per substrate I being 0.746, while in trial 2 the median grams of substrate I used was 0.011, and 0.073 substrate II, with the median proportion of substrate II per substrate I being 0.849.

### Relative humidity did not influence any measured nest trait

In our experiment, ants did not change any wall characteristics with environmental relative humidity levels (linear mixed effects models: wall weight - p = 0.772; Fig 3a, S2 Table; wall length - p = 0.459; Fig 3b, S3 Table, wall area - p = 0.978; Fig 4c, S4 Table, wall density - p = 0.653; Fig 4d, S5 Table, wall substrate composition - p = 0.248; Fig 4e, S6 Table, internal nest area - p = 0.215; Fig 4f, S7 Table). ‘Humidity’ as a fixed effect explained little to none of the variation in the data, while random effects did explain most of the data variation (S2-6 Tables), except for internal nest area (S7 Table). Since we did not find an effect of humidity on any of our measured nest traits, but our sample sizes may be argued to be small, we ran post hoc power analyses to determine the statistical power of each linear mixed effects model. Our models had 80%-85.0% power (therefore at least conventional power) [38] to find any effects of 0.5 (mean difference, β, divided by standard deviation) or higher and 98%-100% power to find any effects of 0.8 or higher, indicating that our negative results are likely not a consequence of low power (S8 Table), and we can conclude that if any effects existed they are likely to be small.

**Fig 3.**
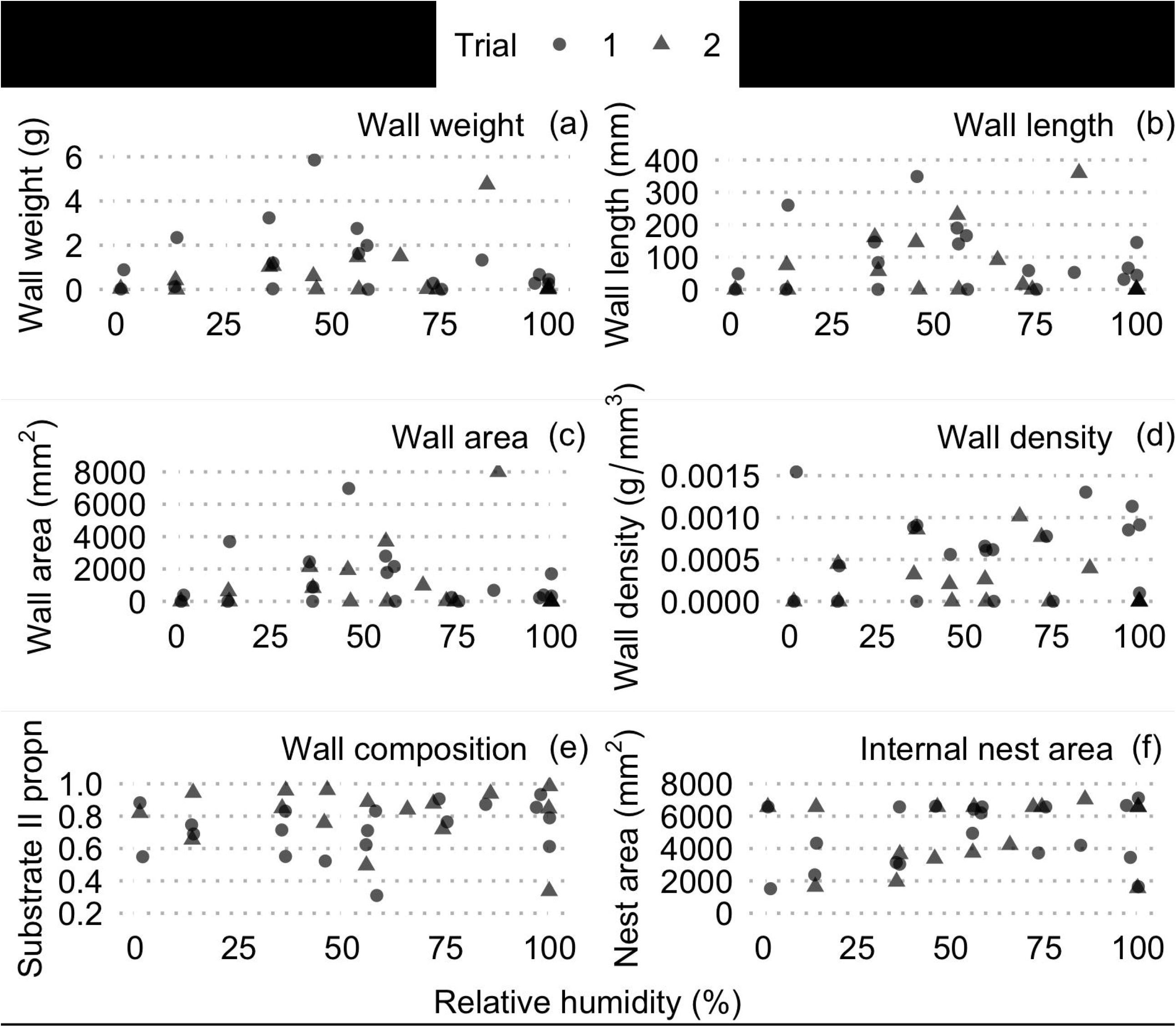
The built nest wall traits measured show no relationship with external humidity levels for trial 1 or 2. Wall weight (a), length (b), area (c), density (d), proportion of substrate II in nest wall (e), and internal nest area (f). Here, and below, trial 1 data points are circles and trial 2 are triangles.

**Fig 4.**
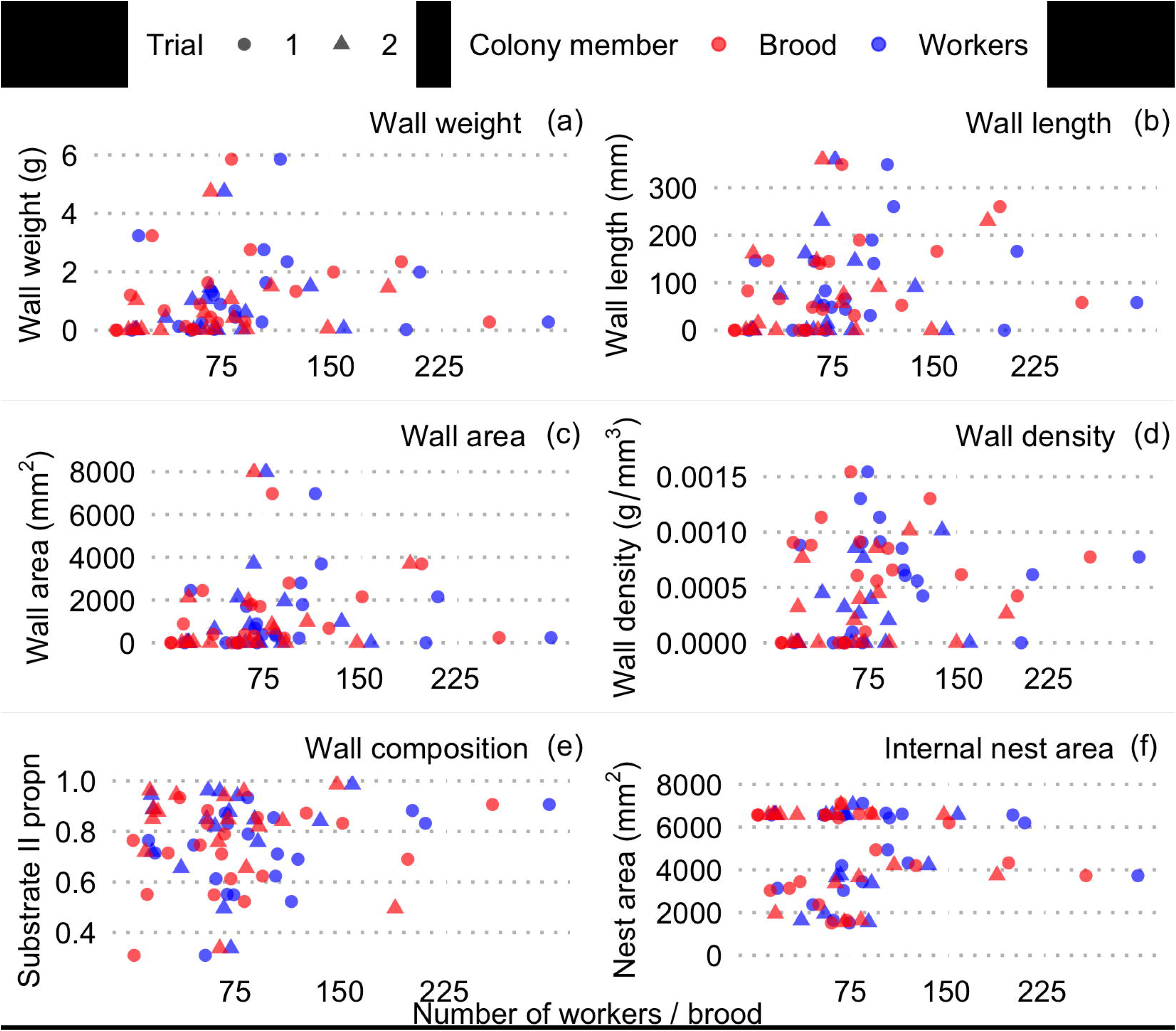
The built nest wall traits measured show no relationship with the number of brood (blue) or workers (red) in a colony for trial 1 or 2. Wall weight (a), length (b), area (c), density (d), proportion of substrate II in nest wall (e), and internal nest area (f).

### No evidence for colony size influencing nest traits

Colonies used in our experiment varied in their demography (note that colony size was calculated by averaging observations of workers and brood from days 1, 5, and 10 of the experiment): the median colony size was 84 workers in trial 1 (range 13-300; 65.3 brood items, range 2-259; 2 queens, range 1-11) and 65.5 workers in trial 2 (range 15-159; 65.7 brood items, range 11 - 190; 2 queens, range 1-11). In our experiment, ant colony size (number of brood or workers) did not influence any wall characteristics (linear mixed effects models: wall weight - brood: p = 0.275, workers: p = 0.517; Fig 4a, S9 Table, wall length - brood: p = 0.054, workers: p = 0.367; Fig 4b, S10 Table, wall area - brood: p = 0.178, workers: p = 0.625; Fig 4c, S11 Table, wall density - brood: p = 0.288, workers: p = 0.448; Fig 4d, S12 Table; wall substrate composition - brood: p = 0.622, workers: p = 0.142; Fig 4e, S13 Table, internal nest area - brood: p = 0.488, workers: p = 0.730; Fig 4f, S14 Table). Since we did not find an effect of colony size on any of our measured nest traits, but our sample sizes may be argued to be small, we ran post hoc power analyses to determine the statistical power of each linear mixed effects model. Our models had 64%-81% power to find any effects of 0.5 (mean difference, β, divided by standard deviation) or higher and 99%-100% power to find any effects of 0.8 or higher, indicating that our negative results are likely not a consequence of low power (S15 Table). Notably, in [25] nest wall area increasing with brood number with an effect size (calculated as we did above) of 0.52 and our model testing for this relationship had 78% statistical power, so we likely had sufficient power to detect an analogous effect.

### Colony mortality did not relate to relative humidity, but larger colonies had higher mortality

We found that worker and brood mortality was highly correlated (β = 1.042 ± 0.168, p < 0.001; S1 Fig, S16 Table). We however found no relationship between relative humidity and the proportion of colony member deaths between trials (10-days) in our generalized linear models (Workers: p = 0.694; Brood: p = 0.756; S2 Fig, S17 Table). We lastly found that the slopes in our linear models predicting colony size in trial 2 from trial 1 were less than 1 (S2 Fig, S18 Table) and were significantly different than models with a slope of 1 (ANOVA: Brood: F = 8.47, p = 0.021; Workers: F = 6.79, p = 0.011; S19 Table). Slopes smaller than 1 indicate that mortality was lower in larger colonies, but since there was no correlation with humidity, the cause is likely not desiccation from low relative humidity or disease (e.g. fungal infection) from high relative humidity.

### Colonies preferred the more porous building substrate, which resembled natural nest walls

We found that ants prefer a more porous substrate: substrate I, the one with smaller grains and thus lower porosity, was not preferred; and in fact, the walls built in our experiment had similar porosity to natural walls collected in the field (Z = 154, p = 0.077), as well as being similar to pure substrate II (Z = 134, p = 0.345) (Fig 5, S20 Table). However, the natural wall substrate was likely not as compact in our porosity assays as in nature, which could possibly change the results. In natural walls, substrate appears tamped down, whereas it was loosely shaken into the test vial for our porosity assays.

**Fig 5.**
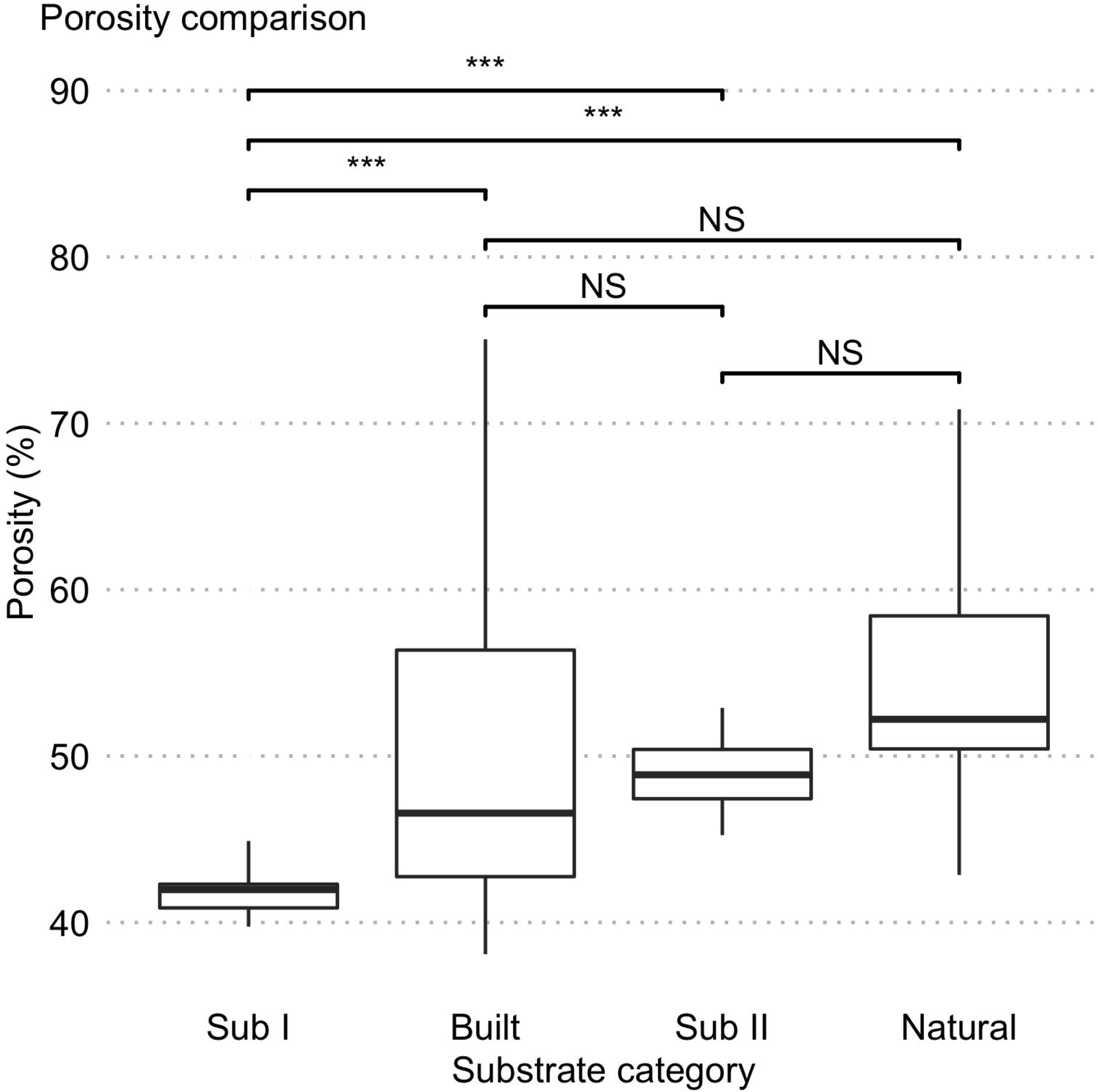
Comparison of porosity between the artificial and natural *Temnothorax rugatulus* nest wall substrates. Bars denote sample medians, boxes represent the first and third quantiles, lines denote the data range. Stars denote significant Dunn’s pairwise multiple comparisons tests.

## Discussion

Our study does not support the hypothesis that environmental humidity affects nest building in *Temnothorax rugatulus*. We found no influence of external humidity on any measured nest property (Fig 3). This might indicate that the nest wall properties are not under selection for maintaining a certain humidity level inside the nest space, or that if they are, they are not plastic (ants do not flexibly adapt them to changing needs for insulation from the environment). We also did not find the effect reported in [25], that nest wall area scales up with brood number in a colony (Fig 4), nor did any other nest property increase with colony size (Fig 4). This inconsistency could stem from different experimental designs – the authors in [25] considered low and high humidity while our study had nine humidity levels - or from the low statistical power of our study. We additionally did not find a relationship between the first relative humidity level that colonies experienced in our study and worker or brood mortality; we did find that larger colonies had overall higher relative mortality (S1-2 Figs). We lastly saw that both our small experimental substrate (substrate II) and the built experimental walls are significantly more porous than the larger experimental substrate but had similar porosity to natural *T. rugatulus* built nest walls. This may be because ants are aiming for porous walls, or it may result from a preference for carrying larger sand grains (which may make the building process more efficient) [23].

Humidity regulation is only one function of social insect nests; however, the internal humidity of the nest can be so important that social insects build structural modifications towards its strict regulation. Examples of these structures include ventilation turrets and thatched nests in *Atta* leafcutter ants [11, 12, 39, 40] and thicker, reinforced mounds along the east-to-west nest axes of *Macrotermes* termites towards retaining water in the nest [13-15]. Here we show in contrast that environmental humidity does not influence *Temnothorax rugatulus* nest wall structure inside nest cavities, suggesting that humidity regulation is not a function of *Temnothorax* nests. Indeed, intrinsic (genetics) rather than extrinsic (temperature and humidity) factors are shown to be more influential towards the nest architectures that harvester ants build [41]. In addition, our *T. rugatulus* colonies selected a less energetically expensive substrate to build with (substrate II), consistent with previous work on wall building substrate choice in *Temnothorax albipennis* colonies [22]. Changing built nest walls in response to humidity might be energetically expensive, which may constrain the flexibility that *T. rugatulus* has in regulating in-nest humidity through wall composition. Additionally, in the desert, external humidity and temperature can change quickly and thus plastic adjustment of nest walls may not be possible, leading to ants building a nest wall structure that is suitable at all levels of humidity. Alternatively, humidity may not be regulated in *T. rugatulus* nests, but instead are physiologically resistant to varying environmental humidity. Instead, *T. rugatulus* colonies may consider other purposes such as nest defense or regulating worker interactions through changing nest densities.

The innate internal humidity of nest cavities may be an important consideration for *Temnothorax* nest site selection. *Temnothorax* ants demonstrate extensive decision-making in house-hunting [42-46], which relates several properties in potential nest cavities. For example, emigrating colonies determine nest size through interactions with other exploring nest mates (quorum sensing) [47]. Also, nest cavities that have smaller nest entrances [44, 48] are more sought after by *Temnothorax* ants for properties such as less light invasion [48]. Additionally, rock-dwelling *Temnothorax albipennis* colonies have been shown to remove substrate from new nest cavities in relation to worker density in the nest (i.e. nest ‘molting’) [27], posing an alternative mechanism that may also regulate in-nest humidity. Therefore, either innate nest cavity properties, or alternative mechanisms to nest wall building, may produce a desirable humidity inside of the nest space (i.e. rock crevices), where *Temnothorax* ants do not need to build nest walls towards its regulation.

Our study is also the first to consider the porosity of the natural and artificial substrates used by *Temnothorax* ants for wall building. Porosity, i.e. the amount of void space between the packed substrate, may influence nest properties in a variety of ways, including thermoregulation and moisture retention, as well as costs of building per volume of wall, none of which has been well studied so far in *Temnothorax* ants. In addition, the soil available to Temnothorax ants in nature likely exhibits a variety of properties that don’t exist in grains of sand, such as the ability to retain moisture in extremely small soil and cellulose grain sizes. The natural nest walls of Temnothorax rugatulus ants can be very densely packed which translated to very slow water penetration in our porosity assays when compared to the virtually instantaneous water penetration of our experimental walls. The mud-brick-like natural *T. rugatulus* nest walls (GC personal observation, but also see Fig 1) may therefore trap humidity differently than loosely packed stone walls. A separate experiment would test this by providing substrates with a variety of weights, types, and sizes could allow rock dwelling ants to select material that produces more compact nest walls than previously possible in both ours and past studies [22, 23, 25]. Alternatively, *Temnothorax* ants may just build with what is available and produce walls with a random mix of substrates that are more energetically efficient to build and those that retain more moisture. Notably, colonies of the ponerine ant *Rhytidoponera metallica* also dwell in rocks and build nest modifications from environmental substrates [49, 50], and other rock dwelling ants differ in their nest size preference [50]. We suggest that exploring the traits of substrates that rock-dwelling ants build nest walls with will provide greater insight to the purpose of these nest walls.

## Acknowledgements

We thank Dr. Nicholas DiRienzo for his insights and help with the photo analysis.

## Supporting information

**S1 Fig. Percentage of brood and worker death in each colony after trial 1 shows no relationship with external humidity levels**.

**S2 Fig. Relatively fewer brood and workers died in larger colonies during the experiment**. Points are the average brood or workers, black lines denote a slope of 1 and red lines are slopes derived from linear models (e.g., formula: log(BroodTrial2) ∼ log(BroodTrial1)).

**S1 Table. Predicted and empirical levels of relative humidity (%) produced from saturated salt solutions**. Predicted relative humidity levels (mean ± standard deviation) are in 20 - 25°C and empirical in 20.40°C ± 0.18°C. Note that magnesium acetate was not used in trial 2 because it produced an inconsistent RH % in trial 1.

**S2 Table. Relationship between built nest wall weight (g) and relative humidity (%)**. Linear mixed effects model: WallWt ∼ Humidity + (1 | Trial)

**S3 Table. Relationship between built nest wall length (mm) and relative humidity (%) for each experimental trial**. Linear mixed effects model: Length ∼ Humidity + (1 | Trial)

**S4 Table. Relationship between built nest wall area (mm**^**2**^**) and relative humidity (%) for each experimental trial**. Linear mixed effects model: Area ∼ Humidity + (1 | Trial)

**S5 Table. Relationship between built nest wall density (g/mm**^**3**^**) and relative humidity (%) for each experimental trial**. Linear mixed effects model: Density ∼ Humidity + (1 | Trial)

**S6 Table. Relationship between the wall substrate composition (proportion of substrate II in build walls) and relative humidity (%) for each experimental trial**. Linear mixed effects model: PropIIWall ∼ Humidity + (1 | Trial)

**S7 Table. Relationship between the internal nest area (mm**^**2**^**) and relative humidity (%) for each experimental trial**. Linear mixed effects model: Nest.Area ∼ Humidity + (1 | Trial)

**S8 Table. Statistical power analyses of linear mixed effects models assessing the effect of humidity on nest properties**. Power analyses considered a moderate (Cohen’s d = 0.5) and high (Cohen’s d = 0.8) effect size. The direction of the effect was taken from the corresponding linear mixed effects model.

**S9 Table. Relationship between built nest wall weight (g) and colony size (number of brood and workers)**. Linear mixed effects model: CollWallWt ∼ Number.Colony + (1 | Trial)

**S10 Table. Relationship between built nest wall length (mm) and colony size (number of brood and workers)**. Linear mixed effects model: Length ∼ Number.Colony + (1 | Trial) **S11 Table. Relationship between built nest wall area (mm**^**2**^**) and colony size (number of brood and workers)**. Linear mixed effects model: Area ∼ Number.Colony + (1 | Trial)

**S12 Table. Relationship between built nest wall density (g/mm**^**3**^**) and colony size (number of brood and workers)**. Linear mixed effects model: Density ∼ Number.Colony + (1 | Trial) **S13 Table. Relationship between built nest wall composition (proportion of substrate II) and colony size (number of brood and workers)**. Linear mixed effects model: PropIIWall ∼ Number.Colony + (1 | Trial)

**S14 Table. Relationship between internal nest area (mm**^**2**^**) and colony size (number of brood and workers)**. Linear mixed effects model: Nest.Area ∼ Number.Colony + (1 | Trial)

**S15 Table. Statistical power of our linear mixed effects models assessing the effect of colony size on nest properties**. Power analyses considered a moderate (Cohen’s d = 0.5) and high (Cohen’s d = 0.8) effect size. The direction of the effect was taken from the corresponding linear mixed effects model.

**S16 Table. Relationship between brood and worker mortality (%) after trial 1**. Linear regression: Brood.Death ∼ Worker.Death

**S17 Table. Relationship between the percentage of worker and brood death in a colony (%) and relative humidity (%)**. The percent death was taken after the first experimental trial. Generalized linear model: worker or brood death ∼ Humidity, family = Binomial

**S18 Table. Relationship between the log of average brood or workers in trials 1 and 2**. Linear regression: Model 1 = Formula: log(avg.worker Trial 2) ∼ log(avg.worker Trial 1); Model 2 = Formula: log(avg.brood Trial2) ∼ 1+ offset(log(avg.brood Trial1))

**S19 Table. Comparing linear models from Table S18 to models with an intercept of 1**. Model comparison: ANOVA(Model 1 ∼ Model 2); Model 1 = Formula: log(avg.worker Trial 2) ∼log(avg.worker Trial 1); Model 2 = Formula: log(avg.brood Trial2) ∼ 1+ offset(log(avg.brood Trial1))

**S20 Table. Comparing the artificial and natural wall substrates porosities (%)**. Dunn’s pairwise tests for wall substrate types: Porosity ∼ SubstrateType

