## Supplemental Table 1 for "*Temnothorax rugatulus* ants do not change their nest walls in response to environmental humidity"

| **Saturated Salt Solution** | **Pred. RH %** | **Emp. RH %** | **Reference** |
| --- | --- | --- | --- |
| *Trial 1* |  |  |  |
| Phosphorous pentoxide (P_2_O_5_) | < 2 x 10^-3^ | 5.79 ± 11.88 | Winston and Bates (1960) |
| Lithium chloride (LiCl) | 12.0 - 12.5 | 13.80 ± 0.36 | Winston and Bates (1960) |
| Magnesium chloride (MgCl) | 32.5 - 33.0 | 36.24 ± 0.52 | Winston and Bates (1960) |
| Potassium carbonate (K_2_CO_3_) | 43.0 - 44.0 | 45.86 ± 0.79 | Winston and Bates (1960) |
| Magnesium nitrate (MgNO_3_) | 53.0 - 55.0 | 55.89 ± 0.73 | Winston and Bates (1960) |
| Magnesium acetate Mg(C_2_H_3_O_2_)_2_ | 65 at 20°C | 58.29 ± 15.34 | Winston and Bates (1960) |
| Sodium chloride (NaCl) | 75.5 - 76.0 | 64.14 ± 11.73 | Winston and Bates (1960) |
| Potassium chloride (KCl) | 83.5 - 84.5 | 78.61 ± 16.35 | Greenspan (1977) |
| Ammonium chloride (NH_4_Cl) | 93.0 - 93.0 | 99.98 ± 0.146 | Winston and Bates (1960) |
| Potassium sulfate (K_2_SO_4_) | 97.5 - 98.0 | 97.35 ± 1.07 | Winston and Bates (1960) |
| *Trial 2* |  |  |  |
| Phosphorous pentoxide (P_2_O_5_) | < 2 x 10^-3^ | 1.08 ± 0.40 | Winston and Bates (1960) |
| Lithium chloride (LiCl) | 12.0 - 12.5 | 13.84 ± 0.33 | Winston and Bates (1960) |
| Magnesium chloride (MgCl) | 32.5 - 33.0 | 35.89 ± 0.69 | Winston and Bates (1960) |
| Potassium carbonate (K_2_CO_3_) | 43.0 - 44.0 | 45.98 ± 0.65 | Winston and Bates (1960) |
| Magnesium nitrate (MgNO_3_) | 53.0 - 55.0 | 55.96 ± 0.48 | Winston and Bates (1960) |
| Sodium chloride (NaCl) | 75.5 - 76.0 | 73.12 ± 1.38 | Winston and Bates (1960) |
| Potassium chloride (KCl) | 83.5 - 84.5 | 86.27 ± 1.02 | Greenspan (1977) |
| Ammonium chloride (NH_4_Cl) | 93.0 - 93.0 | 91.34 ± 19.62 | Winston and Bates (1960) |
| Potassium sulfate (K_2_SO_4_) | 97.5 - 98.0 | 99.86 ± 0.56 | Winston and Bates (1960) |
