## Supplemental Table 2 for "*Temnothorax rugatulus* ants do not change their nest walls in response to environmental humidity"

| Random effects | Name | Variance | Std. dev |  |  |
| --- | --- | --- | --- | --- | --- |
| Trial | (Intercept) | 0.031 | 0.175 |  |  |
| Residual |  | 1.958 | 1.400 |  |  |
| Fixed effects: | β | SE | df | t | p |
| (Intercept) | 1.099 | 0.496 | 9.769 | 2.215 | 0.052 |
| Humidity | -0.002 | 0.008 | 32.033 | -0.293 | 0.772 |
| Model Statistics: |  |  |  |  |  |
| Number of obs: Trial 1 = 19; Trial 2 = 16; | | | | | |
| Marginal R^2^ = 0.002, Conditional R^2^ = 0.018 | | | | | |
