## Supplemental Table 3 for "*Temnothorax rugatulus* ants do not change their nest walls in response to environmental humidity"

| Random effects | Name | Variance | Std. dev |  |  |
| --- | --- | --- | --- | --- | --- |
| Trial | (Intercept) | 7443 | 86.27 |  |  |
| Residual |  | 2755 | 52.49 |  |  |
| Fixed effects: | β | SE | df | t | p |
| (Intercept) | 93.394 | 28.567 | 32.279 | 3.269 | **0.003** |
| Humidity | -0.255 | 0.337 | 18.628 | -0.757 | 0.459 |
| Model Statistics: |  |  |  |  |  |
| Number of obs: Trial 1 = 19; Trial 2 = 16 | | | | | |
| Marginal R^2^ = 0.000, Conditional R^2^ = 0.000 | | | | | |
