## Supplemental Table 4 for "*Temnothorax rugatulus* ants do not change their nest walls in response to environmental humidity"

| Random effects | Name | Variance | Std. dev |  |  |
| --- | --- | --- | --- | --- | --- |
| Trial | (Intercept) | 0 | 0 |  |  |
| Residual |  | 3757310 | 1938 |  |  |
| Fixed effects: | β | SE | df | t | p |
| (Intercept) | 1242.127 | 665.125 | 33 | 1.868 | 0.071 |
| Humidity | -0.291 | 10.433 | 33 | -0.028 | 0.978 |
| Model Statistics: |  |  |  |  |  |
| Number of obs: Trial 1 = 19; Trial 2 = 16 | | | | | |
| Marginal R^2^ = 0.000, Conditional R^2^ = 0.000 | | | | | |
