## Supplemental Table 5 for "*Temnothorax rugatulus* ants do not change their nest walls in response to environmental humidity"

| Random effects | Name | Variance | Std. dev |  |  |
| --- | --- | --- | --- | --- | --- |
| Trial | (Intercept) | 4.308E-08 | 2.076E-4 |  |  |
| Residual |  | 1.836E-07 | 4.285E-04 |  |  |
| Fixed effects: | β | SE | df | t | p |
| (Intercept) | 3.751E-04 | 2.079E-04 | 2.563 | 1.804 | 0.184 |
| Humidity | 1.048E-06 | 2.307E-06 | 32.008 | 0.454 | 0.653 |
| Model Statistics: |  |  |  |  |  |
| Number of obs: Trial 1 = 19; Trial 2 = 16 | | | | | |
| Marginal R^2^ = 0.004, Conditional R^2^ = 0.194 | | | | | |
