## Supplemental Table 6 for "*Temnothorax rugatulus* ants do not change their nest walls in response to environmental humidity"

| Random effects | Name | Variance | Std. dev |  |  |
| --- | --- | --- | --- | --- | --- |
| Trial | (Intercept) | 0.002 | 0.043 |  |  |
| Residual |  | 0.030 | 0.172 |  |  |
| Fixed effects: | β | SE | df | t | p |
| (Intercept) | 0.741 | 0.066 | 5.796 | 11.177 | **<0.001** |
| Humidity | 0.000 | 0.001 | 32.020 | 0.403 | 0.690 |
| Model Statistics: |  |  |  |  |  |
| Number of obs: Trial 1 = 19; Trial 2 = 16 | | | | | |
| Marginal R^2^ = 0.004, Conditional R^2^ = 0.062 | | | | | |
