## Supplemental Table 7 for "*Temnothorax rugatulus* ants do not change their nest walls in response to environmental humidity"

| Random effects | Name | Variance | Std. dev |  |  |
| --- | --- | --- | --- | --- | --- |
| Trial | (Intercept) | 0 | 0 |  |  |
| Residual |  | 37619810 | 1940 |  |  |
| Fixed effects: | β | SE | df | t | p |
| (Intercept) | 4279.503 | 665.538 | 33 | 6.430 | **<0.001** |
| Humidity | 11.149 | 10.439 | 33 | 1.068 | 0.293 |
| Model Statistics: |  |  |  |  |  |
| Number of obs: Trial 1 = 19; Trial 2 = 16 | | | | | |
| Marginal R^2^ = 0.032, Conditional R^2^ = 0.032 | | | | | |
