## Supplemental Table 8 for "*Temnothorax rugatulus* ants do not change their nest walls in response to environmental humidity"

| **Nest property** | β | CI _low_ | CI _high_ | Power |
| --- | --- | --- | --- | --- |
| *Moderate effect* |  |  |  |  |
| Wall weight | -0.022 | 74.87 | 86.19 | 81.00 |
| Wall length | -1.6 | 79.28 | 89.65 | 85.00 |
| Wall area | -31 | 73.78 | 85.31 | 80.00 |
| Wall density | 6.8E-06 | 74.32 | 85.75 | 80.00 |
| Wall composition | 2.7E-03 | 76.51 | 87.50 | 82.50 |
| Internal nest area | 31 | 77.06 | 87.93 | 83.00 |
| *High effect* |  |  |  |  |
| Wall weight | -0.036 | 97.25 | 99.99 | 99.50 |
| Wall length | -2.6 | 96.43 | 99.88 | 99.00 |
| Wall area | -49 | 98.17 | 100.00 | 100.00 |
| Wall density | 1.1E-05 | 97.25 | 99.99 | 99.50 |
| Wall composition | 4.4E-03 | 96.43 | 99.88 | 99.00 |
| Internal nest area | 49 | 97.25 | 99.99 | 99.50 |
| Test Statistics: |  |  |  |  |
| Number of obs: 35; Simulations: 200; alpha = 0.05 | | | | |
