## Supplemental Table 9 for "*Temnothorax rugatulus* ants do not change their nest walls in response to environmental humidity"

| Random effects | Name | Variance | Std. dev |  |  |
| --- | --- | --- | --- | --- | --- |
| *Brood* |  |  |  |  |  |
| Trial | (Intercept) | 0.010 | 0.098 |  |  |
| Residual |  | 1.903 | 1.379 |  |  |
| *Workers* |  |  |  |  |  |
| Trial | (Intercept) | 0 | 0 |  |  |
| Residual |  | 10134 | 100.7 |  |  |
| Fixed effects: | β | SE | df | t | p |
| *Brood* |  |  |  |  |  |
| (Intercept) | 0.651 | 0.383 | 5.569 | 1.701 | 0.144 |
| Number.Colony | 0.004 | 0.004 | 32.535 | 1.110 | 0.275 |
| *Workers* |  |  |  |  |  |
| (Intercept) | 0.758 | 0.416 | 33 | 1.823 | 0.077 |
| Number.Colony | 0.003 | 0.004 | 33 | 0.655 | 0.517 |
| Model Statistics: |  |  |  |  |  |
| Number of obs: Trial 1 = 19; Trial 2 = 16 | | | | | |
| Marginal R^2^: Brood = 0.035; Workers = 0.032 | | | | | |
| Conditional R^2^: Brood = 0.040; Workers = 0.032 | | | | | |
