## Supplemental Table 10 for "*Temnothorax rugatulus* ants do not change their nest walls in response to environmental humidity"

| Random effects | Name | Variance | Std. dev |  |  |
| --- | --- | --- | --- | --- | --- |
| *Brood* |  |  |  |  |  |
| Trial | (Intercept) | 0 | 0 |  |  |
| Residual |  | 9268 | 96.27 |  |  |
| *Workers* |  |  |  |  |  |
| Trial | (Intercept) | 0 | 0 |  |  |
| Residual |  | 1.955 | 1.398 |  |  |
| Fixed effects: | β | SE | df | t | p |
| *Brood* |  |  |  |  |  |
| (Intercept) | 42.085 | 26.286 | 33 | 1.601 | 0.119 |
| Number.Colony | 0.559 | 0.280 | 33 | 1.999 | 0.054 |
| *Workers* |  |  |  |  |  |
| (Intercept) | 60.860 | 29.923 | 33 | 2.034 | 0.050 |
| Number.Colony | 0.264 | 0.289 | 33 | 0.914 | 0.367 |
| Model Statistics: |  |  |  |  |  |
| Number of obs: Trial 1 = 19; Trial 2 = 16 | | | | | |
| Marginal R^2^: Brood = 0.105; Workers = 0.024 | | | | | |
| Conditional R^2^: Brood = 0.105; Workers = 0.024 | | | | | |
