## Supplemental Table 11 for "*Temnothorax rugatulus* ants do not change their nest walls in response to environmental humidity"

| Random effects | Name | Variance | Std. dev |  |  |
| --- | --- | --- | --- | --- | --- |
| *Brood* |  |  |  |  |  |
| Trial | (Intercept) | 0 | 0.098 |  |  |
| Residual |  | 3553566 | 1885 |  |  |
| *Workers* |  |  |  |  |  |
| Trial | (Intercept) | 0 | 0 |  |  |
| Residual |  | 3729954 | 1931 |  |  |
| Fixed effects: | β | SE | df | t | p |
| *Brood* |  |  |  |  |  |
| (Intercept) | 669.848 | 514.723 | 33 | 1.301 | 0.202 |
| Number.Colony | 7.531 | 5.474 | 33 | 1.376 | 0.178 |
| *Workers* |  |  |  |  |  |
| (Intercept) | 993.316 | 574.082 | 33 | 1.730 | 0.093 |
| Number.Colony | 2.732 | 5.544 | 33 | 0.493 | 0.625 |
| Model Statistics: |  |  |  |  |  |
| Number of obs: Trial 1 = 19; Trial 2 = 16 | | | | | |
| Marginal R^2^: Brood = 0.053; Workers = 0.007 | | | | | |
| Conditional R^2^: Brood = 0.053; Workers = 0.007 | | | | | |
