## Supplemental Table 12 for "*Temnothorax rugatulus* ants do not change their nest walls in response to environmental humidity"

| Random effects | Name | Variance | Std. dev |  |  |
| --- | --- | --- | --- | --- | --- |
| *Brood* |  |  |  |  |  |
| Trial | (Intercept) | 3.782E-08 | 1.945E-04 |  |  |
| Residual |  | 1.789E-07 | 4.230E-04 |  |  |
| *Workers* |  |  |  |  |  |
| Trial | (Intercept) | 3.363E-08 | 1.834E-04 |  |  |
| Residual |  | 1.826E-07 | 4.274E-04 |  |  |
| Fixed effects: | β | SE | df | t | p |
| *Brood* |  |  |  |  |  |
| (Intercept) | 3.358E-04 | 1.796E-04 | 1.783 | 1.869 | 0.218 |
| Number.Colony | 1.332E-06 | 1.233E-06 | 32.141 | 1.080 | 0.288 |
| *Workers* |  |  |  |  |  |
| (Intercept) | 3.522E-04 | 1.827E-04 | 2.156 | 1.928 | 0.184 |
| Number.Colony | 9.708E-07 | 1.264E-06 | 32.871 | 0.768 | 0.448 |
| Model Statistics: |  |  |  |  |  |
| Number of obs: Trial 1 = 19; Trial 2 = 16 | | | | | |
| Marginal R^2^: Brood = 0.028; Workers = 0.015 | | | | | |
| Conditional R^2^: Brood = 0.197; Workers = 0.168 | | | | | |
