## Supplemental Table 13 for "*Temnothorax rugatulus* ants do not change their nest walls in response to environmental humidity"

| Random effects | Name | Variance | Std. dev |  |  |
| --- | --- | --- | --- | --- | --- |
| *Brood* |  |  |  |  |  |
| Trial | (Intercept) | 0.002 | 0.046 |  |  |
| Residual |  | 0.029 | 0.171 |  |  |
| *Workers* |  |  |  |  |  |
| Trial | (Intercept) | 0.004 | 0.066 |  |  |
| Residual |  | 0.027 | 0.165 |  |  |
| Fixed effects: | β | SE | df | t | p |
| *Brood* |  |  |  |  |  |
| (Intercept) | 0.743 | 0.057 | 2.824 | 13.010 | **0.001** |
| Number.Colony | 0.000 | 0.000 | 32.280 | 0.498 | 0.622 |
| *Workers* |  |  |  |  |  |
| (Intercept) | 0.700 | 0.068 | 2.290 | 10.225 | **0.006** |
| Number.Colony | 0.001 | 0.000 | 32.915 | 1.505 | 0.142 |
| Model Statistics: |  |  |  |  |  |
| Number of obs: Trial 1 = 19; Trial 2 = 16 | | | | | |
| Marginal R^2^: Brood = 0.007; Workers = 0.057 | | | | | |
| Conditional R^2^: Brood = 0.074; Workers = 0.188 | | | | | |
