## Supplemental Table 14 for "*Temnothorax rugatulus* ants do not change their nest walls in response to environmental humidity"

| Random effects | Name | Variance | Std. dev |  |  |
| --- | --- | --- | --- | --- | --- |
| *Brood* |  |  |  |  |  |
| Trial | (Intercept) | 0 | 0 |  |  |
| Residual |  | 3834729 | 1958 |  |  |
| *Workers* |  |  |  |  |  |
| Trial | (Intercept) | 0 | 0 |  |  |
| Residual |  | 387750 | 1969 |  |  |
| Fixed effects: | β | SE | df | t | p |
| *Brood* |  |  |  |  |  |
| (Intercept) | 5192.885 | 534.699 | 33 | 9.712 | **<0.001** |
| Number.Colony | -3.992 | 5.686 | 33 | -0.702 | 0.488 |
| *Workers* |  |  |  |  |  |
| (Intercept) | 4730.348 | 585.345 | 33 | 8.081 | 0.000 |
| Number.Colony | 1.969 | 5.653 | 33 | 0.348 | 0.730 |
| Model Statistics: |  |  |  |  |  |
| Number of obs: Trial 1 = 19; Trial 2 = 16 | | | | | |
| Marginal R^2^: Brood = 0.014; Workers = 0.004 | | | | | |
| Conditional R^2^: Brood = 0.014; Workers = 0.004 | | | | | |
