## Supplemental Table 15 for "*Temnothorax rugatulus* ants do not change their nest walls in response to environmental humidity"

| **Nest property** | β | CI _low_ | | CI _high_ | | Power ^d^ |
| --- | --- | --- | --- | --- | --- | --- |
| **Brood** |  |  | |  | |  |
| *Moderate effect* |  |  | |  | |  |
| Wall weight | 0.012 | 75.96 | | 87.06 | | 82.00 |
| Wall length | 0.83 | 74.32 | | 85.75 | | 80.50 |
| Wall area | 16 | 71.08 | | 83.09 | | 77.50 |
| Wall density | 3.6E-06 | 74.87 | | 86.19 | | 81.00 |
| Wall composition | 1.5E-03 | 72.15 | | 83.98 | | 78.50 |
| Internal nest area | 17 | 79.50 | | 84/87 | | 79.50 |
| *High effect* |  |  | |  | |  |
| Wall weight | 0.019 | 97.25 | | 99.99 | | 99.50 |
| Wall length | 1.3 | 98.17 | | 100.00 | | 100.00 |
| Wall area | 26 | 97.25 | | 99.99 | | 99.50 |
| Wall density | 5.8E-06 | 98.17 | | 100.00 | | 100.00 |
| Wall composition | 2.4E-03 | 98.17 | | 100.00 | | 100.00 |
| Internal nest area | 27 | 97.25 | | 99.99 | | 99.50 |
| **Workers** |  |  | |  | |  |
| *Moderate effect* |  |  | |  | |  |
| Wall weight | 0.012 | 56.93 | | 70.65 | | 64.00 |
| Wall length | 0.85 | 70.54 | | 82.64 | | 77.00 |
| Wall area | 16 | 67.34 | | 79.93 | | 74.00 |
| Wall density | 3.7E-06 | 74.32 | | 85.75 | | 80.50 |
| Wall composition | 1.4E-03 | 71.61 | | 83.54 | | 78.00 |
| Internal nest area | 17 | 71.08 | | 83.09 | | 77.50 |
| *High effect* |  |  | |  | |  |
| Wall weight | 0.019 | 94.96 | | 99.45 | | 98.00 |
| Wall length | 1.4 | 95.68 | | 99.69 | | 98.50 |
| Wall area | 26 | 95.68 | | 99.69 | | 98.50 |
| Wall density | 6.0E-06 | 97.25 | | 99.99 | | 99.50 |
| Wall composition | 2.3E-03 | 97.25 | | 99.99 | | 99.50 |
| Internal nest area | 27 | 93.58 | | 98.89 | | 97.00 |
| Test Statistics: | | |  | |  | |
| Number of obs: 35; Simulations: 200; alpha = 0.05 | | | | | | |
