## Supplemental Table 16 for "*Temnothorax rugatulus* ants do not change their nest walls in response to environmental humidity"

| Fixed effects: | β | SE | t | p |
| --- | --- | --- | --- | --- |
| (Intercept) | -0.110 | 0.084 | -1.320 | 0.205 |
| Humidity | 1.042 | 0.168 | 6.217 | **<0.001** |
| Model Statistics: |  |  |  |  |
| Number of obs: 18 |  |  |  |  |
| Residual standard error = 0.187; Degrees of freedom: 16 | | | | |
| Multiple R^2^ = 0.707; Adjusted R^2^ = 0.689 | | | | |
| F(1, 16) = 38.65; *p* < 0.001 | | | | |
