## Supplemental Table 17 for "*Temnothorax rugatulus* ants do not change their nest walls in response to environmental humidity"

| Coefficients | β | SE | z | p |
| --- | --- | --- | --- | --- |
| *Workers* |  |  |  |  |
| (Intercept) | -0.010 | 0.894 | -0.011 | 0.991 |
| Humidity | -0.006 | 0.015 | -0.393 | 0.694 |
| *Brood* |  |  |  |  |
| (Intercept) | -0.458 | 0.927 | -0.494 | 0.621 |
| Humidity | -0.005 | 0.015 | -0.310 | 0.756 |
| Model statistics: | | | | |
| Number of obs: 18 | | | | |
| Null deviance: | | | | |
| Workers = 6.0189 on 17 df; Brood = 10.505 on 16 df | | | | |
| Residual deviance: | | | | |
| Workers = 5.863 on 16 df; Brood = 10.408 on 16 df | | | | |
| AIC: Workers = 26.147; Brood = 26.699 | | | | |
| Number of Fisher’s scoring iterations: 3 | | | | |
