## Supplemental Table 18 for "*Temnothorax rugatulus* ants do not change their nest walls in response to environmental humidity"

| Fixed effects: | β | SE | t | | p |
| --- | --- | --- | --- | --- | --- |
| *Brood* |  |  |  | |  |
| Model 1 |  |  |  | |  |
| (Intercept) | 2.033 | 0.652 | 3.118 | | **0.008** |
| log(avg.brood Trial 2) | 0.573 | 0.163 | 3.499 | | **0.003** |
| Model 2 |  |  |  | |  |
| (Intercept) | 0.376 | 0.169 | 2.226 | | **0.042** |
| *Workers* |  |  |  | |  |
| Model 1 |  |  |  | |  |
| (Intercept) | 3.191 | 0.934 | 3.402 | | **0.004** |
| log(avg.workers Trial 2) | 0.330 | 0.230 | 1.430 | | 0.175 |
| Model 2 |  |  |  | |  |
| (Intercept) | 0.501 | 0.195 | 2.565 | | **0.022** |
| Model Statistics: |  | | |  | |
| Number of obs: 16 | | | | | |
| *Brood*:  (Model 1): Residual standard error = 0.574; Degrees of freedom: 14  (Model 2): Residual standard error = 0.676; Degrees of freedom: 15  (Model 1): Multiple R^2^ = 0.467; Adjusted R^2^ = 0.429; (Model 2): N/A  F(1, 14) = 12.24; *p* = 0.004 | | | | | |
| *Workers*:  (Model 1): Residual standard error = 0.638; Degrees of freedom: 14  (Model 2): Residual standard error = 0.781; Degrees of freedom: 15  (Model 1): Multiple R^2^ = 0.128; Adjusted R^2^ = 0.065; (Model 2) N/A  F(1, 14) = 2.045; *p* = 0.175 | | | | | |
