## Supplemental Table 19 for "*Temnothorax rugatulus* ants do not change their nest walls in response to environmental humidity"

| Model: | Res. Df | RSS | Df | | | Sum of Sq | | F | | p |
| --- | --- | --- | --- | --- | --- | --- | --- | --- | --- | --- |
| *Brood* | 15 | 9.14 | -1 | | | -3.45 | | 8.47 | | **0.011** |
| *Workers* | 1515 | 6.85 | -1 | | | -2.24 | | 6.79 | | **0.021** |
| Model Statistics: |  | | |  |  | |  | |  | |
| Number of obs: 16 | | | | | | | | | | |
