## Supplemental Table 20 for "*Temnothorax rugatulus* ants do not change their nest walls in response to environmental humidity"

| Substrate comparison: | Z | p |
| --- | --- | --- |
| Substrate I vs. Substrate II | -3.128 | **0.009** |
| Substrate I vs. Natural | 4.102 | **<0.001** |
| Substrate I vs. Built | 2.771 | **0.011** |
| Substrate II vs. Natural | 26 | 0.075 |
| Substrate II vs. Built | 134 | 0.345 |
| Natural vs. Built | 154 | 0.077 |
| Test Statistics: |  |  |
| Number of obs: substrates I, II, and natural = 12; built walls = 10 | | |
| **Kruskall-Wallis test:**  X^2^ = 51, df = 51, p = 0.474 | | |
