## Supplementary figures and images for "*Temnothorax rugatulus* ants do not change their nest walls in response to environmental humidity"

### Supplemental Figure 1

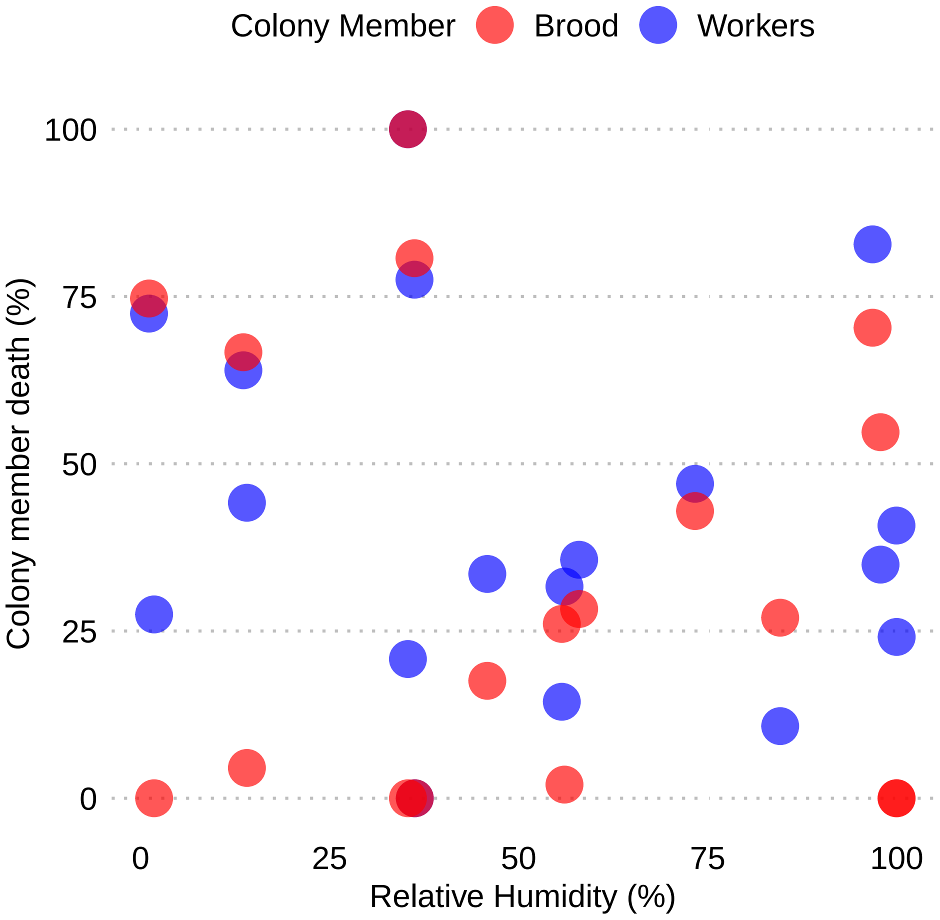

### Supplemental Figure 2

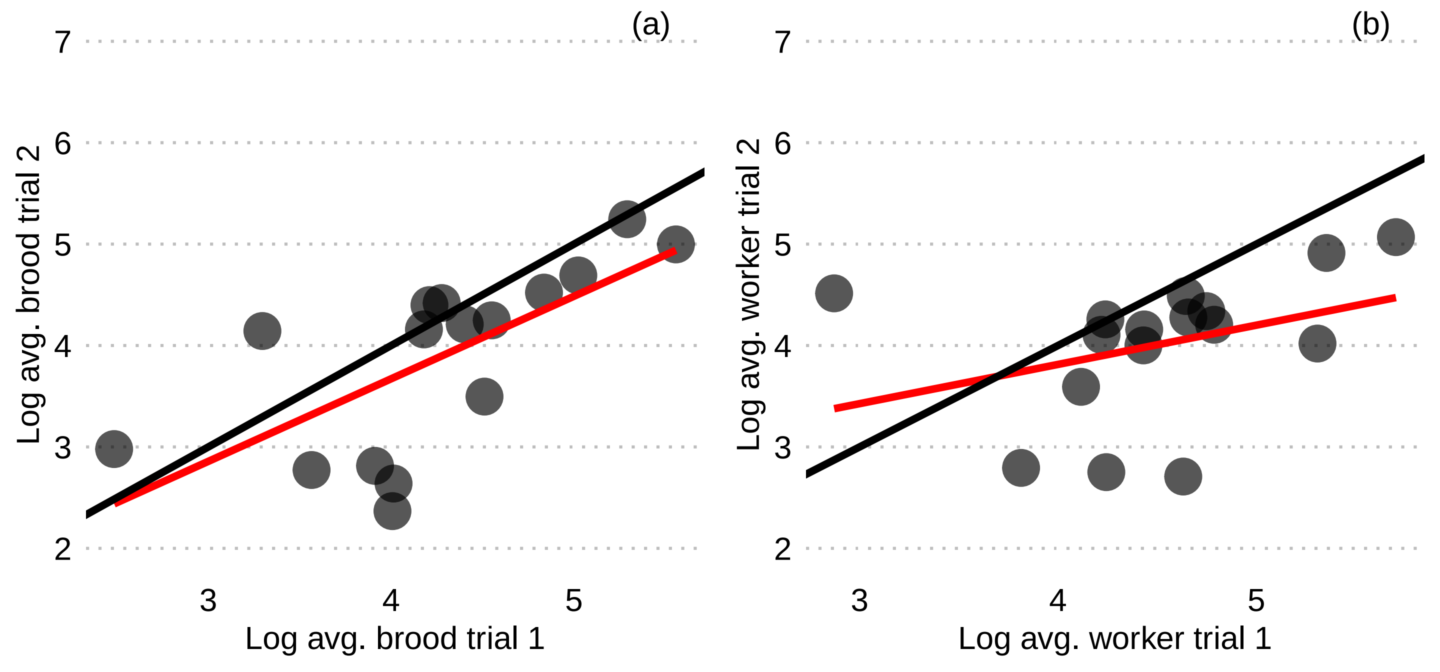
